# Axial asymmetry organizes division plane orthogonality in *Neisseria gonorrhoeae*

**DOI:** 10.1101/2024.06.28.601173

**Authors:** Aditya C. Bandekar, Diego A. Ramirez-Diaz, Samantha G. Palace, Yi Wang, Ethan C. Garner, Yonatan H. Grad

## Abstract

For rod-shaped bacterial model organisms, the division plane is defined by the geometry of the cell. However, for *Neisseria gonorrhoeae*, a coccoid organism that most commonly exists as a diplococcus and that possesses genes coding for rod-based cell division systems, the relationship between cell geometry and division is unclear. Here, we characterized the organization of *N. gonorrhoeae* division using a combination of fluorescent probes, genetics, and time-lapse microscopy. We found that the planes of successive cell divisions are orthogonal and temporally overlapping, thereby maintaining diplococcal morphology. Division takes place perpendicular to a subtle long-axis in each coccus. In keeping with the ParABS and the MinCDE systems reading the long-axis of rod-shaped bacteria, in the coccoid *N. gonorrhoeae*, ParB segregates along this subtle long-axis and cells lacking *minCDE* have severe morphological consequences, including an inability to perform orthogonal division and aberrant assembly of the division plane at the cell poles. Taken together, this stresses the central role of even slight dimensional asymmetry as a general organizational principle in bacterial cell division.

## Introduction

Much of our current understanding of prokaryotic cell division, an important determinant of cell shape, has been elucidated in the rod-shaped organisms (bacilli) *Escherichia coli* and *Bacillus subtilis*. However, bacteria come in a variety of shapes and sizes^1^, including spherical (cocci), ellipsoid (ovococci), curved, spirals, branched, and star-shaped. Since the genes that code for the core cell division proteins are often conserved among bacteria^2^, this raises the question of how this morphological diversity is achieved.

A hallmark of bacterial division is binary fission – the ability to give rise to two equally sized daughter cells. In most bacillary and ovococcal^3^ mother cells, the only division plane that can generate two equally sized daughter cells is at mid-cell, perpendicular to the cell’s long-axis. Notable exceptions are the gut symbiont *Laxus oneistus* and oral symbionts of the *Neisseriaceae* family that divide longitudinally^4–6^. Regardless of whether the bacillus divides perpendicular or parallel to the long-axis, in rod-shaped cells, successive division planes are always parallel to each other.

In coccoids, an infinite number of possible division planes could divide the cell into equal halves. *Staphylococcus aureus*, a Gram-positive coccus, divides in alternating, perpendicular planes^7,8^. Spherically shaped mutants of *E. coli* also employ orthogonal successive division planes^9,10^. *Neisseria gonorrhoeae* is a Gram-negative organism that is primarily observed as a diplococcus^11,12^, raising several questions: how does *N. gonorrhoeae* choose its division plane, how does it maintain diplococcal morphology through divisions, and how does its individual coccal and diplococcal morphology influence the geometry of division?

Transmission electron microscopy of thin sections of dividing gonococci^13,14^ and phase contrast microscopy of live cells^14^ suggest that successive division planes are perpendicular to each other. However, studying the underlying molecular mechanisms of *N. gonorrhoeae* division at a finer scale with conventional light microscopy has proved elusive due to *N. gonorrhoeae*’s relatively small size (600 nm – 800 nm), fastidious growth, and the limited genetic tools with which to create fluorescent reporter strains.

Many systems work together to ensure that bacterial division occurs in the right place and at the right time after the chromosome has been duplicated and segregated into the future daughter cells. This segregation needs to occur prior to cytokinesis to avoid the formation of anucleate cells and the chromosome being trapped in the division septa. Thus, systems including ParABS^15^, Muk^16^ and SMC^17^ segregate DNA, and proteins including SlmA^18^ and Noc^19,20^ prevent division plane assembly over the nucleoid. Cell geometry plays an important role in chromosome segregation^21^. Theoretical models^22^ and experimental work^23^ have shown that ParA/ParB systems move the origins of DNA replication along the long-axis of the cell^24–26^ by ParB moving along a gradient of ParA bound to the nucleoid.

In addition to chromosome segregation, protein gradient systems like MinCDE play crucial roles in determining division site placement^27,28^ by oscillating along the cell’s long-axis in *E. coli*^29,30^. Modeling the behavior of this system in spherical cells suggests that a mere 5% difference in the length between the long and short axes is sufficient for the Min system to begin oscillating^30–32^. In *N. gonorrhoeae*, *minCDE* plays a role in maintaining cell integrity^33^. While MinD’s role in division site placement has not been studied *in situ*, fluorescent *N. gonorrhoeae* MinD heterologously expressed in *E. coli* oscillates along the long-axis^34^.

In this study, we first investigated the organization of successive division planes in *N. gonorrhoeae* and their relationship to its diplococcal morphology. Additionally, since *N. gonorrhoeae* has *parABS* and *minCDE* genes, we investigated whether subtle axial asymmetry offered by the coccoid shape played a role in organizing division plane orientation in *N. gonorrhoeae*.

## Results

### Successive division planes are perpendicular in *N. gonorrhoeae*

To investigate the role of division plane selection in generating diplococci, we generated a fluorescent reporter strain (nAB019) of *N. gonorrhoeae* in which we fused the green fluorescent protein mNeonGreen (mNG)^35^ to the N-terminus of the cell division protein ZapA^36^ using a modified version (See Methods) of an allelic exchange system^37^. This fusion appeared to maintain ZapA function because it localized to the division site at mid-cell, manifesting as a straight line or a ring depending on the orientation of the cell relative to the viewing plane (**Figure 1A, 1B**). Time-lapse microscopy of nAB019 revealed that the division plane rotated orthogonally every generation (**Figure 1C, Supplementary Video 1, Supplementary Video 2, Supplementary Video 3**). Due to the cells being under an agarose pad during imaging, divided cells remained in close proximity to each other, making visualization of single cells difficult after microcolonies reached ∼ 8 cells (two successive divisions). Similarly, we could not observe the separation of daughter diplococci from parental diplococci.

**Figure 1.**
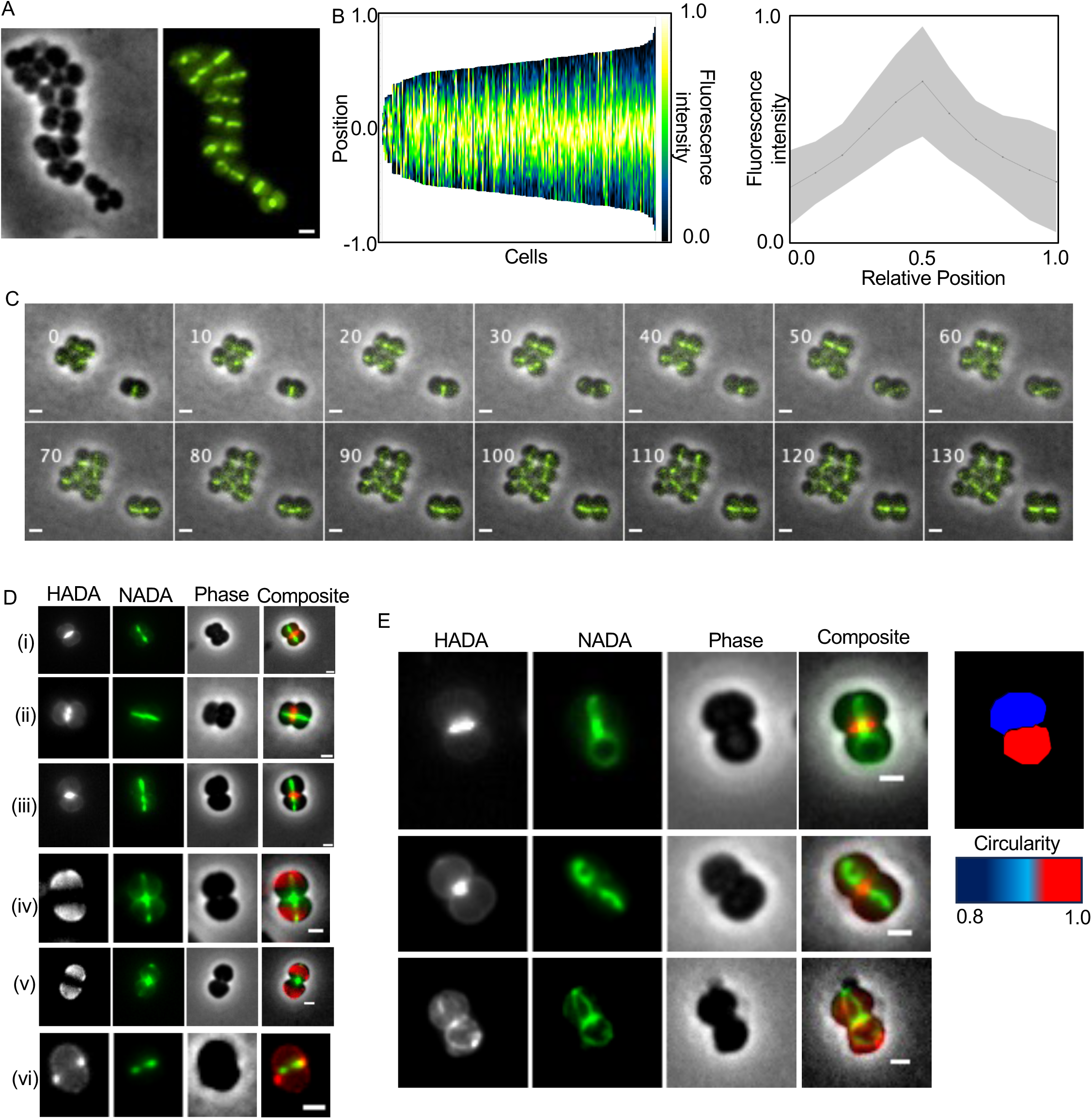
Successive division planes are orthogonal in *N. gonorrhoeae.* All scale bars are 1 μm. (A) Localization pattern of mNeonGreen-ZapA in single cells of *N. gonorrhoeae* strain nAB019. (L) Phase contrast image. (R) mNeonGreen-ZapA localization. (B) Population kymograph (L) (n=193 cells) and fluorescence intensity profile (R) (n=11 cells) of mNG-ZapA in nAB019. The black line is the mean fluorescence intensity. Gray envelope is standard deviation. (C) Live cell imaging of nAB019. Time of acquisition of each image is indicated in minutes. (D) Examples of *N. gonorrhoeae* cells sequentially labeled with two FDAAs. HADA labeling was first performed for 45 minutes (column 1, grey), cells were washed and then labeled with NADA for 45 minutes (column 2, green). Column 3 shows the phase contrast image. Column 4 shows the composite image (HADA in red, NADA in green). (E) Examples of *N. gonorrhoeae* cells where sister cells have rotated relative to each other. Sequential labelling with two FDAAs was performed as in Figure 1D. Heatmap indicates circularity (1.0 = circular)

To assess the orientation of successive division planes in liquid culture, we next performed fluorescent d-amino acid (FDAA) labeling as markers of new cell wall deposition during cell division^38^. We first labeled with blue-fluorescent HADA (3-[[(7-Hydroxy-2-oxo-2*H*-1-benzopyran-3-yl) carbonyl] amino]-D-alanine) followed by green-fluorescent NADA (3-[(7-Nitro-2,1,3-benzoxadiazol-4-yl) amino]-D-alanine). Similar to **Figure 1C**, FDAA labeling showed that cells built their septa in a perpendicular orientation to their parents **(Figure 1D, (i)-(iii))**. Additionally, since the signal from the division plane stained in the first labeling (HADA) persisted during the labeling of the second division plane (NADA), we concluded that the mother cell was still dividing as the daughter cells began the process of division. In cells at a different cell cycle stage during the labeling, the same temporal overlap can be observed [**Figure 1D, (iv), (v)]:** the second FDAA (NADA) has labeled both division planes, whereas the HADA channel is dark at the first division plane. This indicated the latter occurred because NADA labeled the first division plane while concomitantly labeling the second division plane. This temporal overlap between successive division planes explains how diplococcal morphology is maintained during cell divisions.

These experiments also revealed several other features of cell division. First, division begins at the periphery and subsequently proceeds inward toward the cell interior [**Figure 1D**, **(vi)**]. Second, division planes in sister cells were not always parallel to each other (**Figure 1E**), suggesting that as the septum holding together the two cells of the diplococcus matures, each cell can rotate relative to its sister cell. Third, the top sister cell (**Figure 1E** row 1, false-colored blue) is less circular than the bottom sister cell (false-colored red), indicating that *N. gonorrhoeae* cells are not perfectly circular, containing subtle dimensional asymmetry with a long/short axis ratio of ∼1.2 (1.253 ± 0.15) (**Figure 2A**), which became oriented differently in these sister cells during the imaging process.

**Figure 2.**
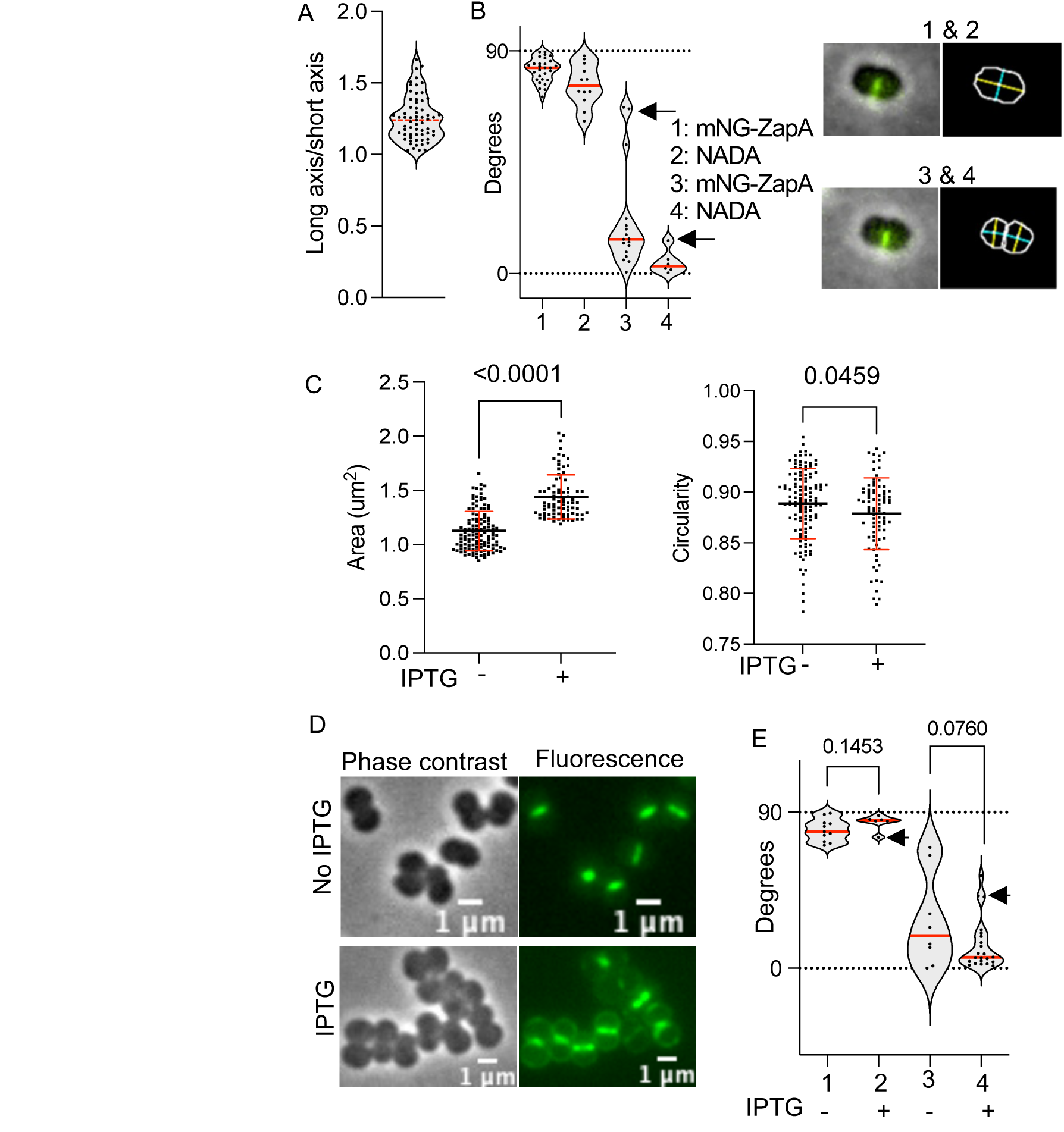
The division plane is perpendicular to the cellular long-axis. All scale bars are 1 μm. (A) Long-axis: short axis ratio of Cellpose segmented masks of phase contrast images. n=63 cells. Red line is the mean long axis :short axis ratio. (B) Angle between the long-axis and the current generation’s division plane (1,2) and the previous generation’s division plane (3,4). 1 and 3 are from measuring mNG-ZapA in nAB019. 2 and 4 use data from *N. gonorrhoeae* labelled with NADA. Arrows indicate subset of cells where the sister cells have rotated relative to each other. Inset (top row) shows a representative cell used for calculating 1 and 2. Inset (bottom row) shows a representative cell used for calculating 3 and 4. Inset (L) shows a composite image (phase contrast in gray, mNG-ZapA in green). Inset (R) shows a Cellpose generated mask. Overlayed on this mask is a yellow line/s indicating the long-axis and a teal line/s indicating the short axis. (C) Characteristics of GCGS0457 cells expressing a lac-inducible copy of PBP1. Left-No induction. Right-Induction with 0.5mM IPTG for 2.5 hours. p-values of unpaired t-test with Welch’s correction are indicated. (D) Phase contrast and green fluorescence images of NADA labelled *N. gonorrhoeae* expressing a lac-inducible copy of PBP1. Top row-no induction. Bottom row PBP1 induction with 0.5mM IPTG after 2.5 hours. (E) Angle between the long-axis and the current generation division plane (1,2) or the previous generation division plane (3,4) in NADA labelled cells expressing normal levels of PBP1 (1 and 3) or overexpressing PBP1 using 0.5mM IPTG for 2.5 hours (2 and 4). P-values of unpaired t-test with Welch’s correction are indicated. Arrows indicate subset of cells where the sister cells have rotated relative to each other.

### The division plane is perpendicular to the cellular long-axis

To investigate the role of cell shape in setting the orientation of division planes, we next measured the orientation of the long-axis relative to the division plane. Cells were segmented using Cellpose^39^; the long-axis of the cell masks was oriented nearly perpendicular to the current division plane [**Figure 2B (1)-**mNG-ZapA: 82.3° ± 4.8°, **Figure 2B (2)-**NADA: 76° ± 8.3°] and nearly parallel to the mother cell division plane [**Figure 2B (3)-**mNG-ZapA: 20.7° ± 20.6°; **Figure 2B (4)-**NADA: 4.5° ± 4.7°].

To test whether the orthogonal relationship between the division plane and the long-axis holds even in cells with morphologies that deviate from wild-type cells, we overexpressed the peptidoglycan assembly enzyme, *ponA* (*pbp1*) (**Supplementary** Figure 1). As the FDAA signal is restricted to the septum in wild-type *N. gonorrhoeae* (**Figure 1D**), we reasoned that by overexpressing *ponA*, we might drive the cell to deposit cell wall along the periphery. Doing so would alter the aspect ratio of the cell, thereby increasing the long-axis: short-axis ratio.

Consistent with this hypothesis, we found that relative to the area of wild-type cells (1.125 ± 0.18 µm^2^, n=121 cells), cells overexpressing PBP1 are larger (1.441 ± 0.2 µm^2^, n=83 cells, unpaired t-test p<0.0001) and less circular (0.8786 ± 0.03 versus 0.8886 ± 0.03 for wild type cells, unpaired t-test p=0.0459) (**Figure 2C**), with NADA signal observed around the cell periphery as well as the septum (**Figure 2D**). Despite these perturbations, the angle between the division plane and long-axis (84.19° ± 4.4°) was similar (unpaired t-test, p=0.1453) to cells without PBP1 overexpression (79.7° ± 6.3°) (**Figure 2E**). Taken together, these data indicate the division plane is oriented roughly perpendicular to the cell’s long-axis.

### The ParABS system segregates chromosomes along the long-axis of *N. gonorrhoeae*

Next, we determined the localization of the ParABS system, which reads the long-axis of the cell. We hypothesized that if a physiologically relevant long-axis exists in *N. gonorrhoeae*, ParB should segregate along it. We N-terminally tagged ParB with the red fluorescent protein mScarlet3^40^ in nAB019 to generate strain nAB055. The ParB in this fusion appeared functional because it localized to regions of the cell where the nucleoid was expected (**Figure 3A**). As seen in other ParA/B systems, imaging revealed that the two ParB foci mirrored each other as cells grew, localizing to 0.25 and 0.75 lengths of the cell (**Figure 3B**). In some cells, >2 ParB foci were observed, suggesting the presence of multiple copies of the chromosome (**Figure 3C**). Unlike the division plane, the axis of segregation of ParB foci occurred nearly parallel to the long-axis, with an angle of ≤20° in ∼85% of cells (**Figure 3D**). In sister cells that moved relative to each other, the axis of ParB segregation was parallel to the long-axis of its cell, even if it was not parallel to the division plane of the mother cell (**Figure 3E**). Interestingly, when we tracked the segregation of ParB foci and ZapA localization in nAB055 using live cell microscopy, ParB foci segregated (**Figure 3F-** red box) before ZapA was first observed at the division site (**Figure 3F-** green box) (**Supplementary Video 4, Supplementary Video 5**), suggesting that new long axes are established long before division occurs. To test this, we observed the growth of nAB019. As cells progressed through their cell cycle, cell constriction occurred at the division site (**Figure 3G**, box 1). This constriction created new long axes in the subsequent daughter cells, which were perpendicular to the previous axis (**Figure 3G**, box 2). The next division plane was then placed perpendicular to this long-axis (**Figure 3G**, box 3).

**Figure 3.**
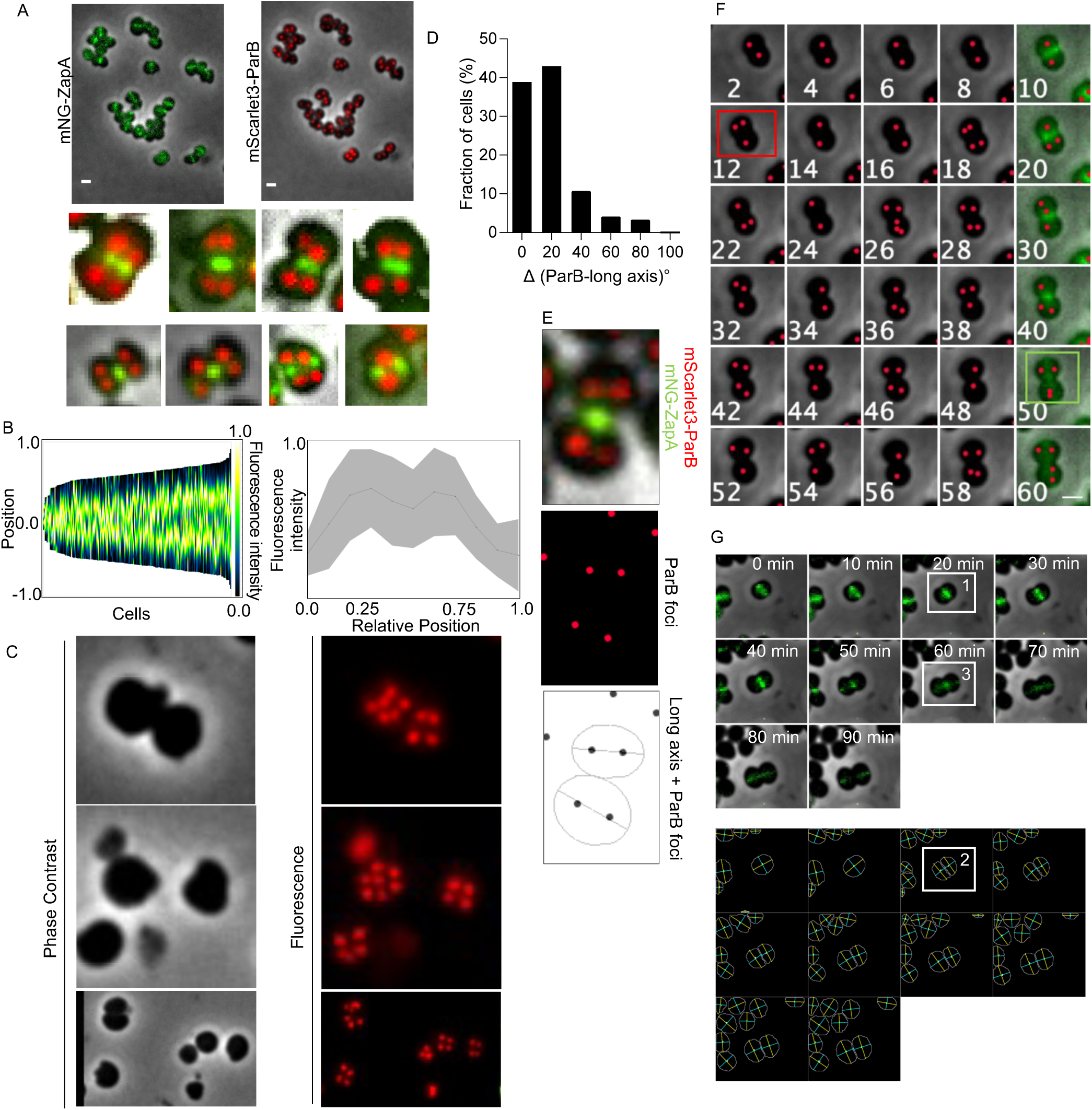
The ParABS systems segregates chromosomes along the long-axis. All scale bars are 1 μm. (A) Localization pattern of mNeonGreen-ZapA and mScarlet3-ParB in single cells of *N. gonorrhoeae* nAB055. Insets show examples of zoomed-in cells to show the ParB localization patterns clearly. (B) Population kymograph (L) and fluorescence intensity profile (R) of mScarlet3-ParB in *N. gonorrhoeae* strain nAB055 (n=193 cells). (C) Localization of mScarlet3-ParB in multinucleate *N. gonorrhoeae* strain nAB055. (D) Angle of ParB segregation axis relative to the long-axis of *N. gonorrhoeae* nAB055. (E) Images showing an example of a *N. gonorrhoeae* nAB055 cell rotated relative to its sister cell. (Top) Composite image showing cell boundary, mNeonGreen ZapA localization and mScarlet3-ParB localization. (Middle) ParB foci (red dots) determined using LoG detector in TrackMate. (Bottom) Long-axis of Cellpose generated mask (black line) overlayed with ParB foci (black dots). (F) Montage showing live cell imaging of *N. gonorrhoeae* nAB055. mScarlet3-ParB was imaged every 2 minutes, mNG-ZapA was imaged every 10 minutes. Red dots are ParB foci. Red box indicates the time point when ParB segregation was first detected. Green box indicates the time ZapA was first detected at the assembling division plane. (G) Rotation of division plane in successive generations of *N. gonorrhoeae* nAB019 cells. (Top) Montage of timelapse imaging of nAB019, images taken every 10 minutes. (Bottom) Montage of Cellpose segmented cells. The yellow line is the long-axis. The blue line is the short axis.

Together, these data suggest that as *N. gonorrhoeae* proceeds through its cell cycle, a long-axis develops, along which the segregation of ParB occurs, followed by the assembly of the division plane perpendicular to the cell’s long-axis.

### MinCDE is necessary for the division plane to be perpendicular to the long-axis of *N. gonorrhoeae*

A long/short axis ratio of ∼1.2 [similar to *N. gonorrhoeae* (**Figure 2A**)] was demonstrated to be the minimal ratio for the *min* system to read out the long axis of chambers *in vitro*^32^. As MinCDE is known to oscillate along the long-axis and play a critical role in division site selection, we hypothesized that if we deleted the *minCDE* genes, successive division planes would no longer be orthogonal. We created an unmarked deletion of the *minCDE* operon using allelic exchange in nAB019. As expected, this strain showed severe morphological defects in cell size and shape (**Figure 4A**). Further, its ability to assemble the division ring at mid-cell was severely compromised (**Figure 4B**), resulting in the formation of mini-cells (**Figure 4A**, arrows). *ΔminCDE* cells were unable to rotate their division planes, causing them to no longer be perpendicular to the long-axis (**Figure 4C**) and making cells unable to separate from each other during division (**Figure 4D**). Taken together, these data suggest that the long-axis sensing of MinCDE is necessary for orthogonal division site placement in *N. gonorrhoeae*.

**Figure 4.**
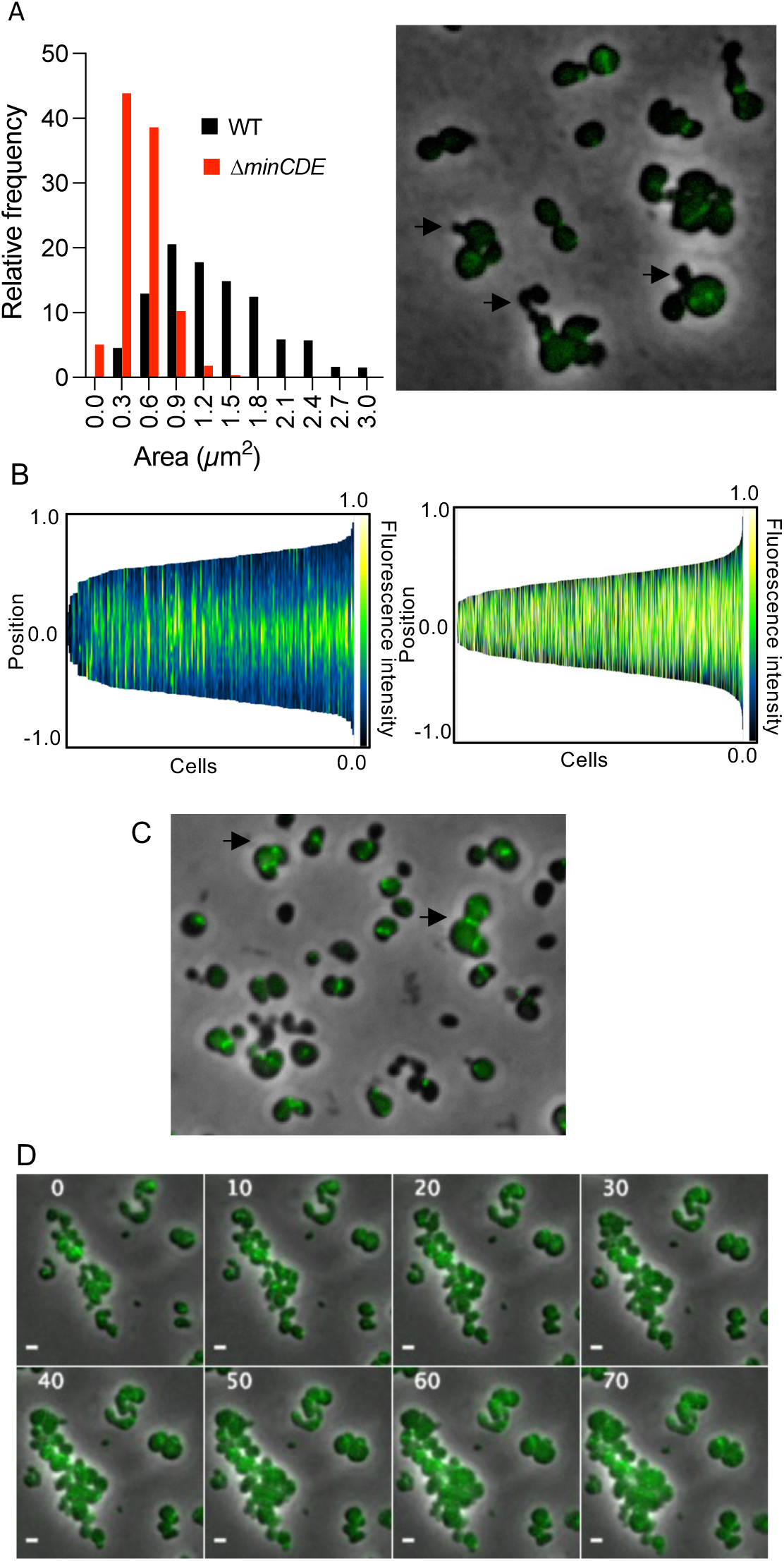
The *minCDE* system is necessary for division plane fidelity. All scale bars are 1 μm. (A) Histograms of cell lengths of nAB019 (L) and nAB019 *ΔminCDE* (R). Fluorescence micrograph showing mini-cell formation (arrows). (B) Population kymograph of mNG-ZapA in nAB019 (L) (n=193 cells) and nAB019 *ΔminCDE* (R) (n=850 cells). (C) Fluorescence micrograph showing mNG-ZapA localization in *N. gonorrhoeae* nAB019 *ΔminCDE* cells where successive division planes are not orthogonal (Arrows). (D) Timelapse imaging of nAB019 *ΔminCDE*. Images were taken every 10 minutes.

## Discussion

This work demonstrates how the diplococcal morphology of *N. gonorrhoeae* is generated and maintained. Our data supports models that have been proposed for the role of axial asymmetry in organizing perpendicular division planes in coccoid bacteria^30,41^. Building on these models, **Figure 5** shows a proposed model for how cell division occurs, specifically in *N. gonorrhoeae*.

**Figure 5.**
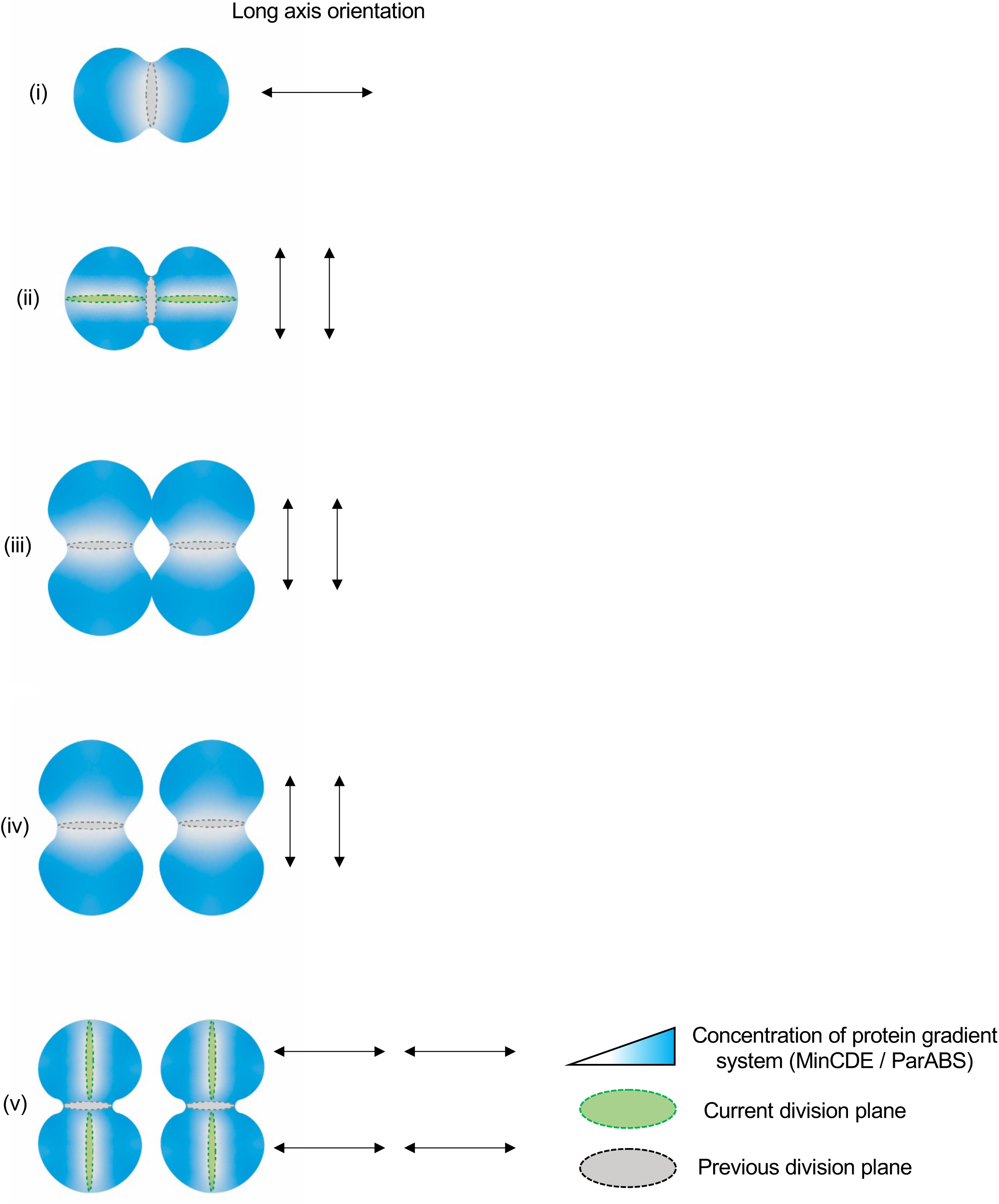
Proposed model of growth and division in *N. gonorrhoeae*. (i) MinCDE reads subtle asymmetry to constrain division perpendicular to the mother cell’s long axis (horizontal black arrow). (ii) Ongoing cytokinesis generates two new vertical long axes in the daughter cells. Protein gradient systems flip to vertical oscillations. A new division plane (gray ring) begins to be assembled perpendicular to the previous division plane (black ring). (iii)-(iv) With the division planes specified, cell cycle proceeds towards cytokinesis. (v) Step (ii) is repeated for the next generation, generating four new horizontal long axes.

First, the MinCDE system reads out small asymmetries in cell shape, oscillating along the long axis [**Figure 5(i)**, horizontal arrow], resulting in MinCDE constraining the first division plane (gray ring) to be perpendicular to the cell’s long-axis. Division plane assembly and the start of cell constriction precedes the formation of two new long axes in the daughter cells which are perpendicular to the long-axis of the mother cell [**Figure 5(ii),** vertical arrows]. The MinCDE system reads these new long axes and now oscillates orthogonally to the previous axis of oscillation. While we did not determine the point during the cell cycle at which the MinCDE and ParABS systems begin to read out the new long axes, we propose this occurs when oscillation is no longer feasible along the long-axis of the mother cell, due to septal closure, impedance of MinDE oscillations by the division plane, and/or some other geometric parameter causing MinDE to prefer oscillating along the new long-axis. New division planes then begins assembly perpendicular to the previous division plane, consistent with observations in *S. aureus*^7,8^ and spherical mutants of *E.coli*^10^. In *N. gonorrhoeae* however, these division planes are temporally overlapping, with a new plane beginning to be assembled (green ring) before the previous plane (gray ring) has been completely resolved **[Figure 5(ii)]**. This overlap in division cycles creates a situation where cells are perpetually in some stage of septation, resulting in diplococci. Once the division planes of the current generation are specified, the cell cycle proceeds toward cytokinesis [**Figure 5(iii-iv)],** culminating with **Figure 5(ii)** being repeated for the grand-daughter cells, creating four new long axes [**Figure 5(v)**, horizontal arrows].

Our data shows the ParABS and MinCDE systems of *N. gonorrhoeae* read the subtle (long-axis: short-axis ratio ∼1.2) axial asymmetry of cells, aligning with prior observations of how these systems sense the long-axis of rod-shaped cells. While this work shows that the subtle long-axis plays a role in *N. gonorrhoeae* division plane orientation, demonstration of a causal role of the long-axis in this process requires the ability to generate and maintain spherical *N. gonorrhoeae* as well as tools to physically manipulate *N. gonorrhoeae* to create a new long-axis.

It is intriguing to speculate about why coccoid bacteria rotate their division planes every generation. We propose that the planes are orthogonal because there is no other option that creates two sisters that are equally sized and inherit an entire genome, since nucleoid segregation occurs parallel to the long-axis. The next question is why nucleoid segregation occurs along the long-axis. We propose that exploiting subtle axial asymmetry to segregate nucleoids and rotate division planes allows *N.* gonorrhoeae, a human pathogen that relies on host cell adherence and immune evasion for its pathogenesis, to maintain its coccoid morphology and size, traits that enhance its ability to colonize mucosal surfaces^42^ and reduce its sensing by the immune system^43,44^.

## Methods

### Culturing Conditions

*N. gonorrhoeae* was cultured as described^45^. Briefly, glycerol stocks *of N. gonorrhoeae* were streaked for single colonies on GCB agar (Difco) with Kellogg’s supplement (GCB-K) and incubated at 37 °C with supplemental 5%CO_2_ for 16-18 hours. Liquid culture was performed in phosphate-buffered gonococcal (GCP) medium (15 g /l proteose peptone 3 (Thermo-Fisher), 1 g /l soluble starch (Thermo-Fisher), 4 g/l K_2_HPO_4_, 1 g /l KH_2_PO_4_, 5 g /l NaCl) supplemented with Kellogg’s supplement (GCP-K) and incubated on an orbital shaker at 200rpm in a 37°C incubator with 5% environmental CO_2_.

### Static and live cell microscopy

Cells were grown as described in “Culturing Conditions” above. 1.2µl of cell suspension was placed in a low evaporation 50 mm glass-bottomed dish (MatTek Corporation - No. 1.5) and a phosphate buffered saline (pH 7.4 Gibco) pad containing 2.5% agarose was placed on top of the cells. For live cell imaging, the pad was made with GCB-K. Epifluorescence and phase images were collected using a Nikon Ti-E inverted, widefield microscope equipped with a Nikon Perfect Focus system, a Piezo Z drive motor, a Nikon Plan Apo λ ×60/×100 1.4NA objective, an Andor Zyla VSC-04459 sCMOS camera, NIS Elements (v4.5) and a stage top incubator (Okolab) set to 37°C (for live cell imaging) and equipped with 5% environmental CO2. To reduce drift due to temperature fluctuations, the sample was mounted on the microscope and allowed to equilibrate to the imaging chamber temperature for 10-15 minutes before image acquisition.

Fluorescence was captured using a 6-channel Spectra X LED light source and a Sedat Quad filter set. The excitation (Ex.) and emission (Em.) filters used in this study were: Ex. 395±25nm and Em. 435±25nm for HADA; Ex. 470±24nm and Em. 515±25nm for green fluorophores (mNeonGreen and NADA); Ex. 550±15nm and Em. 595±25nm for mScarlet3.

### FDAA labelling

Cells were grown as described in “Culturing Conditions” above. The overnight growth was transferred to 10ml of GCP-K to a density of Abs600 0.2. When the Abs600 reached 0.4 (∼ 60 minutes later), 1ml of cell suspension was centrifuged at 4,000g for 1 minute and the cell pellet was resuspended in GCP-K (pre-warmed to 37°C) containing 100µM HADA (generously provided by the Eric Rubin lab). This was incubated on an orbital shaker at 200rpm for 45 minutes in a 37°C incubator with environmental 5%CO2. Next, cells were rapidly washed with 2 ml pre-warmed (37°C) GCP-K twice and transferred to pre-warmed (37°C) GCP-K containing 1mM NADA (Tocris Bioscience; Cat. No. 6648) and incubated in the same conditions as above. Cells were harvested by centrifugation at 4,000g for 1 minute and immediately fixed by resuspending in 70% ice cold ethanol. These stained and fixed cells were harvested by centrifugation at 10,000g for 1 minute, resuspended in 1X Phosphate Buffered Saline (Difco, pH 7.4) and then prepared for static microscopy as described above in “Static and live cell microscopy”.

### Sequences of fluorescent proteins and linker mScarlet3 (codon optimized for N. gonorrhoeae)

ATGGATTCGACTGAGGCTGTAATCAAAGAGTTCATGCGGTTTAAGGTCCATATGGAGGGCAGCATGAATGGCCACGAATTCGAAATAGAAGGCGAAGGCGAAGGTCGTCCTTACGAAGGTACCCAAACGGCAAAACTTCGGGTCACGAAGGGCGGCCCCCTGCCCTTCTCGTGGGACATTCTGAGCCCGCAATTCATGTACGGTTCGCGCGCATTCACCAAGCATCCTGCGGACATCCCTGATTACTGGAAACAGAGCTTTCCCGAGGGTTTCAAATGGGAGCGCGTTATGAACTTCGAAGATGGCGGCGCCGTCAGCGTTGCACAGGATACTTCGCTGGAGGATGGCACCTTGATTTACAAAGTCAAGTTGCGGGGCACAAACTTTCCTCCTGACGGTCCGGTTATGCAGAAAAAGACTATGGGCTGGGAAGCCTCCACAGAGCGGCTTTACCCTGAGGACGTTGTTCTGAAAGGCGACATTAAGATGGCTCTTCGTTTGAAAGACGGTGGTCGGTATCTTGCGGATTTCAAGACAACGTATCGCGCAAAGAAGCCGGTACAGATGCCTGGTGCCTTTAATATCGACCGTAAATTGGATATTACTAGCCACAATGAAGACTATACAGTAGTAGAGCAATATGAGCGCAGCGTGGCTCGGCATTCCACGGGTGGCTCTGGCGGTTCC

### mNeonGreen (codon optimized for N. gonorrhoeae)

ATGGTCAGCAAGGGCGAAGAGGACAACATGGCTTCGTTGCCGGCAACGCACGAGCTGCACATATTCGG CTCGATCAATGGCGTGGATTTTGATATGGTGGGCCAGGGCACGGGCAACCCCAACGATGGTTACGAGG AGCTGAACCTTAAATCCACTAAGGGTGACTTGCAGTTCTCTCCTTGGATATTGGTTCCGCACATCGGTTA CGGCTTTCACCAATACTTGCCGTACCCGGACGGCATGAGCCCTTTTCAAGCGGCAATGGTGGATGGCAG CGGTTACCAGGTTCATCGTACCATGCAGTTTGAGGATGGCGCGAGCCTTACGGTTAATTATCGGTACAC ATATGAGGGCTCGCATATTAAGGGTGAGGCTCAGGTCAAAGGCACAGGCTTTCCGGCTGACGGTCCCG TTATGACCAACAGCCTTACTGCTGCTGACTGGTGTCGGAGCAAAAAGACGTATCCGAATGACAAGACTA TCATAAGCACTTTTAAGTGGAGCTACACGACCGGTAATGGTAAACGTTATCGCTCGACCGCACGCACTAC TTATACATTTGCTAAACCGATGGCGGCGAATTATTTGAAGAACCAGCCTATGTACGTGTTCCGGAAAACT GAACTTAAACACTCCAAAACTGAGCTTAACTTTAAGGAGTGGCAAAAGGCCTTCACAGACGTGATGGGC ATGGATGAATTGTATAAG

### 15aa linker (codon optimized for *N. gonorrhoeae*)

CTCGAGGGCAGCGGTCAGGGCCCTGGCTCCGGCCAAGGTAGCGGC

### Construction of fluorescent reporter strains and the *ΔminCDE* strain

Fluorescent reporter strains and deletion mutants of *N. gonorrhoeae* were generated using a modified version of a published allelic exchange method^37^. Briefly, allelic exchange was achieved in two steps. In step 1, using kanamycin positive selection, an intermediate *N. gonorrhoeae* strain was generated, where the target gene was replaced with a dual selection *aph3-galK* cassette. In step 2, using 2-deoxygalactose (2-DOG) negative selection, the final *N. gonorrhoeae* strain was generated, where the *aph3-galK* cassette was replaced by the target gene fused with the fluorescent tag of choice or deleted in the case of a deletion mutant. **nAB019** [*zapA::ɸ(mNeonGreen-zapA)*]

The intermediate strain nAB008 was generated by transforming FA19 with a plasmid assembled using Gibson Assembly (New England Biolabs) containing 4 fragments: 1) PCR with primers oAB112 and oAB098 and FA19 genomic DNA as template (containing the region upstream of *zapA*); 2) PCR with primers oAB097 and oAB096 and *aph3-galk* cassette as template; 3) PCR with primers oAB095 and oAB113 and FA19 genomic DNA as template (containing the region downstream of *zapA*); 4) PCR with primers oAB114 and oAB115 and *HindIII*-digested pUC19 DNA as template (the vector backbone).

The final strain nAB019 was generated by transforming nAB008 with a plasmid containing 5 fragments: 1) PCR with primers oAB112 and oAB082 and FA19 genomic DNA as template (containing the region upstream of *zapA*); 2) PCR with primers oAB093 and oAB106 and synthetic *mNeonGreen* (IDT; attached to 15aa linker via PCR) as template; 3) PCR with primers oAB107 and oAB110 and FA19 genomic DNA as template (containing *zapA*); 4)PCR with primers oAB105 and oAB113 and FA19 genomic DNA as template (containing the region downstream of *zapA*); 5)PCR with primers oAB114 and oAB115 and *HindIII*-digested pUC19 DNA as template (the vector backbone).

### **nAB055** [zapA::ɸ(mNeonGreen-zapA) parB::ɸ(mScarlet3-parB)]

The intermediate strain was generated by transforming nAB019 with a plasmid assembled using Gibson Assembly (New England Biolabs) containing 4 fragments: 1) PCR with primers oAB128 and oAB130 and FA19 genomic DNA as template (containing the region upstream of *parB*); 2) PCR with oAB132 and oAB131 and *aph3-galk* cassette as template; 3) PCR with primers oAB133 and oAB129 and FA19 genomic DNA as template (containing the region downstream of *parB*); 4) PCR with primers oAB114 and oAB115 and *HindIII*-digested pUC19 DNA as template (the vector backbone).

The final strain nAB055 was generated by transforming the above strain with a plasmid containing 5 fragments: 1) PCR with primers oAB128 and oAB134 and FA19 genomic DNA as template (containing the region upstream of *parB*); 2) PCR with primers oAB136 and oAB135 and synthetic *mScarlet3* (IDT; attached to 15aa linker via PCR) as template; 3) PCR with primers oAB138 and oAB137 and FA19 genomic DNA as template (containing *parB*); 4) PCR with primers oAB139 and oAB129 and FA19 genomic DNA as template (containing the region downstream of *parB*); 5) PCR with primers oAB114 and oAB115 and *HindIII*-digested pUC19 DNA as template (the vector backbone).

### **nAB113** [zapA::ɸ(mNeonGreen-zapA) ΔminCDE]

The intermediate strain nAB094 was generated by transforming nAB019 with a plasmid assembled using Gibson Assembly (New England Biolabs) containing 4 fragments: 1)PCR with primers oAB146 and oAB145 and FA19 genomic DNA as template (containing the region upstream of *minCDE*); 2)PCR with primers oAB148 and oAB147 and *aph3-galk* cassette as template; 3)PCR with primers oAB150 and oAB149 and FA19 genomic DNA as template (containing the region downstream of *minCDE*); 4)PCR with primers oAB114 and oAB115 and *HindIII*-digested pUC19 DNA as template (the vector backbone).

The final strain nAB113 was generated by transforming the above strain with a plasmid containing 3 fragments: 1) PCR with primers oAB146 and oAB151 and FA19 genomic DNA as template (containing the region upstream of *minCDE*); 2)PCR with primers oAB152 and oAB149 and FA19 genomic DNA as template (containing the region downstream of *minCDE*); 3)PCR with primers oAB114 and oAB115 and *HindIII*-digested pUC19 DNA as template (the vector backbone).

The sequences of all plasmids were confirmed using long-read sequencing (Oxford Nanopore Technologies, Plasmidsaurus) before proceeding to the next step. Plasmids were linearized using *XbaI* (New England Biolabs) prior to transformation.

## Oligonucleotides used in this study

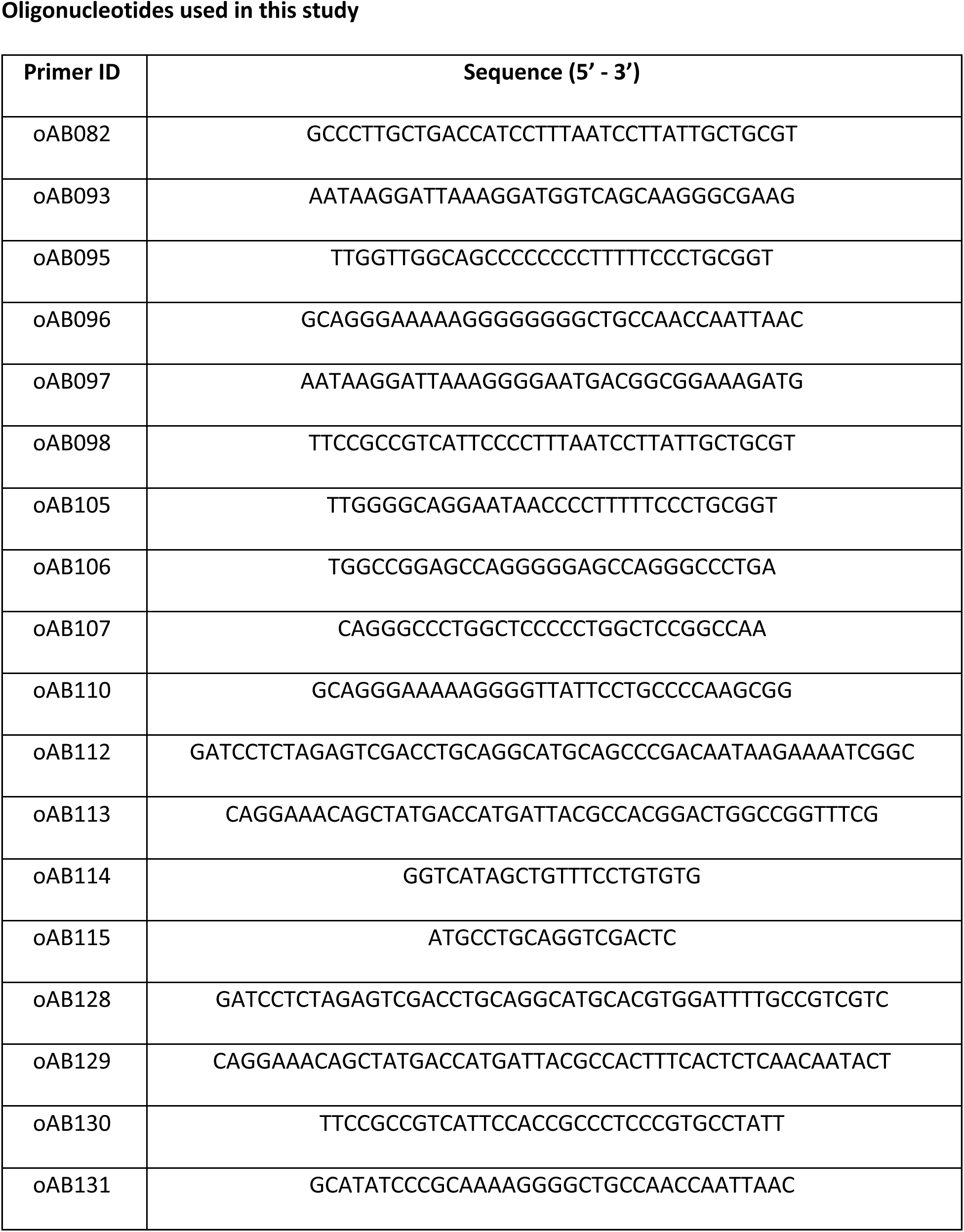
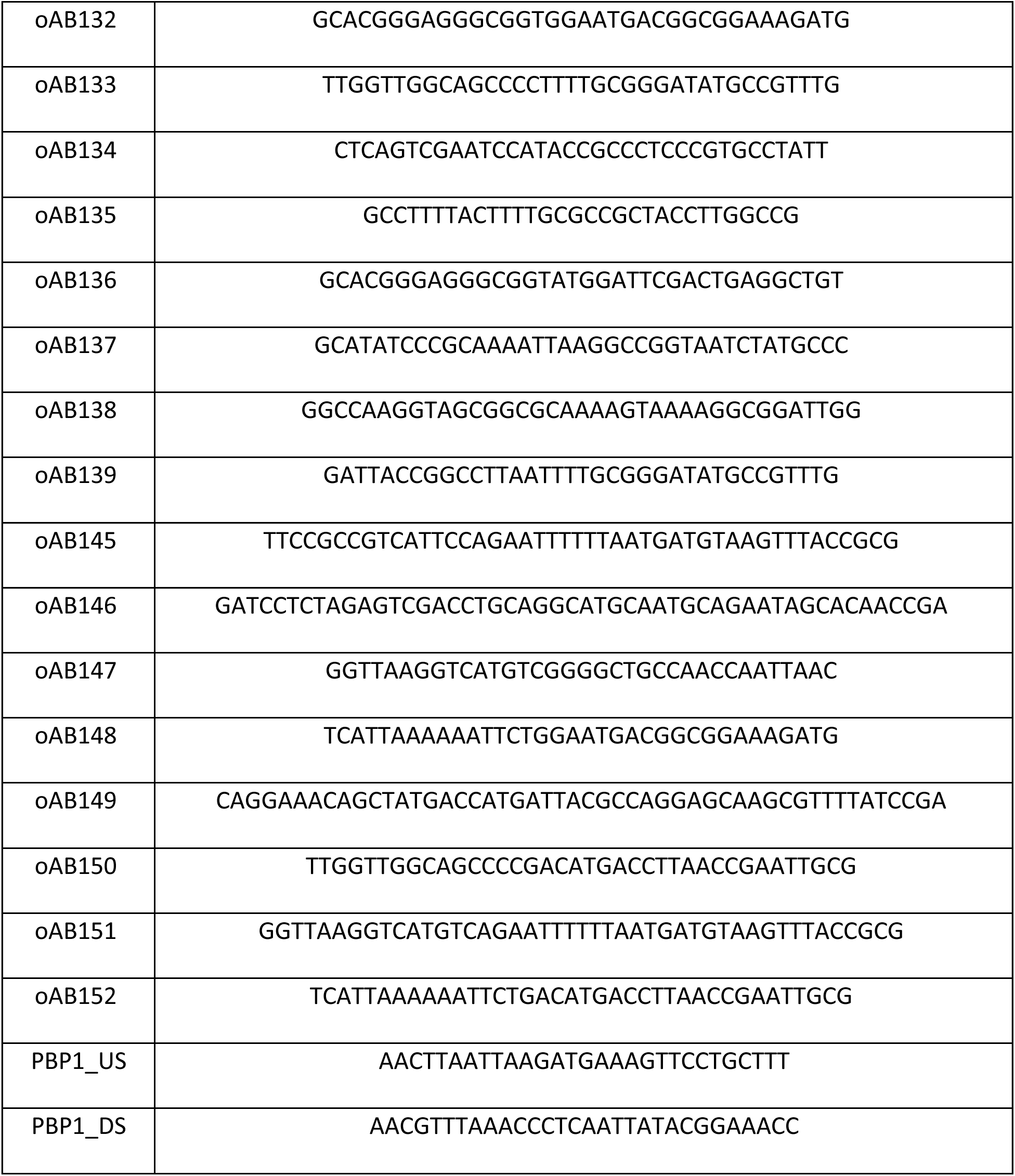

### Transformation of *N. gonorrhoeae*

*N. gonorrhoeae* that was to be transformed was first grown for 16–18 h on GCB-K plates at 37 °C in 5 % CO2. Piliated colonies (20 to 30) were picked and transferred to 150—200µl of GCP-K. Cells (30 µl) were spotted onto GCB-K agar and the spots were allowed to dry. The restriction enzyme digest reaction (containing ∼ 400-800 ng of linearized plasmid DNA) was spotted on top of the dried spot of cells and allowed to dry. CutSmart buffer (New England Biolabs)-only spots were used as a negative control. Plates were incubated for 6-8 h at 37°C with environmental 5%CO2. The growth from the spot was resuspended in 100µl of GCP-K and plated onto GCB-K agar containing either 100µg/ml kanamycin (for step1) or 1% 2-DOG (for step 2) and incubated for 24–36 h at 37 °C with environmental 5%CO2. Step 2 transformants that were confirmed to be kanamycin sensitive and DOG resistant upon sub-culture were assessed by PCR and Sanger sequencing.

## Strains created in this study

| ID | Strain | Genotype |
| --- | --- | --- |
| nAB019 | FA19 | *zapA::φ(mNeonGreen-zapA)* |
| nAB055 | FA19 | *zapA::φ(mNeonGreen-zapA) parB::φ(mScarlet3-parB)* |
| nAB113 | FA19 | *zapA::φ(mNeonGreen-zapA) ΔminCDE* |
| YWC02 | GCGS0457 | *p<sub>lac</sub> ponA* |

### Construction of PBP1 overexpression strain

The effect of PBP1 overexpression was examined in GCGS0457, a ceftriaxone-susceptible clinical isolate that contains the PBP1^L421P^ allele^46^. The *ponA* gene encoding PBP1 was amplified with its native promoter from GCGS0457 using primers PBP1_US and PBP1_DS. Next, it was introduced into the pKH37 complementation vector^47^ between the *PacI* and *PmeI* restriction sites, placing it downstream of the lac inducible promoter. The pKH37::*ponA* plasmid and the empty vector pKH37 control were methylated with *HaeIII* methyltransferase (New England Biolabs), linearized with *PciI* (New England Biolabs), and transformed into GCGS0457 via spot transformation as described^48^. Transformants were selected with 1 μg/mL chloramphenicol. Overexpression of the PBP1 protein was confirmed by bocillin-FL measurement of PBP abundance.

### PBP abundance measurement

Protein abundance of PBP1 was calculated using the fluorescent penicillin derivative bocillin-FL (Thermo Fisher). GCGS0457, GCGS0457(pKH37), and GCGS0457(pKH37::*ponA*) from overnight cultures were suspended in liquid GCP medium (15 g/L protease peptone 3, 1 g/L soluble starch, 4 g/L dibasic K2HPO4, 1 g/L monobasic KH2PO4, 5 g/L NaCl) supplemented with 1% IsoVitalex (Becton Dickinson) and 0.042% NaHCO3 to a density of OD600 0.1. Suspensions were incubated with aeration at 37°C for 2.5-3 hours. Bacterial cells were collected by centrifugation, washed once with 1 mL of sterile phosphate-buffered saline (PBS), and resuspended in PBS with 5 μg/mL bocillin-FL and 0.1% dimethyl sulfoxide (DMSO) to a final concentration of 1 mL of OD600 0.5 per 50 μL suspension. Bocillin-FL suspensions were incubated for 5 minutes. An equal volume 2x SDS-PAGE sample buffer (Novex) was added, and samples were boiled for 5 minutes. Proteins in 30 μL of each suspension were separated by SDS-PAGE on a 4-12% Tris-Glycine protein gel (Novex), which was visualized on a Typhoon imager (Amersham) (excitation 488 nm/emission 526 nm) to detect bocillin-FL fluorescence. Gels were then stained with GelCode™ Blue Stain Reagent (ThermoFisher) and visualized with white light to allow total protein normalization between samples. Densitometry was performed with FIJI^49^.

### Image analysis

All images were analyzed using FIJI. Phase contrast images were used for cell segmentation which was performed using Cellpose^39^. For Figure 2A, 2B, 2E and 3G, the long axes of Cellpose generated masks were determined using an ImageJ macro. Source code can be found at https://github.com/abandekar/Ngo-cell-division/tree/main.

### Calculation of the angle between long-axis and division plane

The angle between the long-axis and the division plane (Figure 2B, Figure 2E) was calculated in FIJI. First, the background fluorescence of the epifluorescence image of mNG-ZapA was subtracted (rolling ball radius = 50 pixels) and then converted to a binary image. Next, each septum was fitted to an ellipse, and the long-axis of this ellipse was determined using the same macro as above. This gave us the angle of the division plane. The angle between the long-axis and the division plane was then calculated. The kymographs in Figure 1B, Figure 3B and Figure 4B were generated using MicrobeJ^50^.

### Calculation of angle between long-axis and ParB foci

First, the background fluorescence of the epifluorescence image of mScarlet3-ParB was subtracted (rolling ball radius = 50 pixels) in FIJI. Next, ParB foci were determined using the Laplacian of Gaussian (LoG) detector in Trackmate^51^ with an estimated object diameter of 0.25 microns. Masks were generated using Cellpose as above. Next, custom Python code (available at: https://github.com/diegoalejandrord/angle_parb) was used to analyze these two images (the ParB foci image and the masks image). This code performed two functions. First, cells were filtered out to retain only those cells which have n=2 ParB foci. Second, the angle between the two ParB foci and the long-axis was determined.

## Acknowledgements

We would like to thank Junhao Zhu from the Eric Rubin laboratory for guidance with microscopy and for generously providing HADA. We thank Robert Nicholas for providing the *N. gonorrhoeae* FA19 strain. We thank Lucia Ricci for assisting with the graphic design of the model presented in Figure 5. This work was funded by the National Institutes of Health grants AI132606 and AI153521 to Y.H.G.

## Author contributions

The work was conceptualized by A.C.B., E.C.G., and Y.H.G. The PBP1 overexpression strain used in Figure 2 was constructed by Y.W. and S.G.P. The code to determine the angle between ParB foci and the long-axis that generated the data for Figure 3D was written by D.A.R. All other experiments and analyses were performed by A.C.B. Original draft was written by A.C.B. The draft was reviewed and edited by A.C.B., E.C.G., and Y.H.G. Funding was acquired by Y.H.G. The work was supervised by E.C.G. and Y.H.G.

## References

1. Young, K. D. The Selective Value of Bacterial Shape. Microbiol Mol Biol Rev 70, 660–703 (2006).

2. Megrian, D., Taib, N., Jaffe, A. L., Banfield, J. F. & Gribaldo, S. Ancient origin and constrained evolution of the division and cell wall gene cluster in Bacteria. Nat Microbiol 7, 2114–2127 (2022).

3. van Raaphorst, R., Kjos, M. & Veening, J.-W. Chromosome segregation drives division site selection in Streptococcus pneumoniae. Proceedings of the NaSonal Academy of Sciences 114, E5959–E5968 (2017).

4. Leisch, N. et al. Growth in width and FtsZ ring longitudinal positioning in a gammaproteobacterial symbiont. Curr Biol 22, R831–832 (2012).

5. Leisch, N. et al. Asynchronous division by non-ring FtsZ in the gammaproteobacterial symbiont of Robbea hypermnestra. Nat Microbiol 2, 16182 (2016).

6. Nyongesa, S. et al. Evolution of longitudinal division in multicellular bacteria of the Neisseriaceae family. Nat Commun 13, 4853 (2022).

7. Monteiro, J. M. et al. Cell shape dynamics during the staphylococcal cell cycle. Nat Commun 6, 8055 (2015).

8. Saraiva, B. M. et al. Reassessment of the distinctive geometry of Staphylococcus aureus cell division. Nature CommunicaSons 11, 4097 (2020).

9. Iwaya, M., Goldman, R., Tipper, D. J., Feingold, B. & Strominger, J. L. Morphology of an Escherichia coli mutant with a temperature-dependent round cell shape. Journal of Bacteriology 136, 1143–1158 (1978).

10. Begg, K. J. & Donachie, W. D. Division Planes Alternate in Spherical Cells of Escherichia coli. Journal of Bacteriology 180, 2564–2567 (1998).

11. Neisser, A. Ueber eine der Gonorrhoe eigentümliche Micrococusform. CentralblaU für die medizinischen WissenschaXen 17, 497–500 (1879).

12. Kampmeier, R. H. Identification of the gonococcus by Albert Neisser. 1879. Sex Transm Dis 5, 71–72 (1978).

13. Fitz-James, P. Thin sections of dividing neisseria gonorrhoeae. Journal of Bacteriology 87, 1477–1482 (1964).

14. Westling-Häggström, B., Elmros, T., Normark, S. & Winblad, B. Growth paàern and cell division in Neisseria gonorrhoeae. Journal of Bacteriology 129, 333–342 (1977).

15. Ogura, T. & Hiraga, S. Partition mechanism of F plasmid: two plasmid gene-encoded products and a cis-acting region are involved in partition. Cell 32, 351–360 (1983).

16. Hiraga, S. et al. Chromosome partitioning in Escherichia coli: novel mutants producing anucleate cells. J Bacteriol 171, 1496–1505 (1989).

17. Briàon, R. A., Lin, D. C. & Grossman, A. D. Characterization of a prokaryotic SMC protein involved in chromosome partitioning. Genes Dev 12, 1254–1259 (1998).

18. Bernhardt, T. G. & de Boer, P. A. J. SlmA, a Nucleoid-Associated, FtsZ Binding Protein Required for Blocking Septal Ring Assembly over Chromosomes in E. coli. Mol Cell 18, 555– 564 (2005).

19. Wu, L. J. & Errington, J. Coordination of cell division and chromosome segregation by a nucleoid occlusion protein in Bacillus subtilis. Cell 117, 915–925 (2004).

20. Veiga, H., Jorge, A. M. & Pinho, M. G. Absence of nucleoid occlusion effector Noc impairs formation of orthogonal FtsZ rings during Staphylococcus aureus cell division. Mol Microbiol 80, 1366–1380 (2011).

21. Wu, L. J. et al. Geometric principles underlying the proliferation of a model cell system. Nat Commun 11, 4149 (2020).

22. Hu, L., Vecchiarelli, A. G., Mizuuchi, K., Neuman, K. C. & Liu, J. Directed and persistent movement arises from mechanochemistry of the ParA/ParB system. Proceedings of the NaSonal Academy of Sciences 112, E7055–E7064 (2015).

23. Vecchiarelli, A. G., Neuman, K. C. & Mizuuchi, K. A propagating ATPase gradient drives transport of surface-confined cellular cargo. Proceedings of the NaSonal Academy of Sciences 111, 4880–4885 (2014).

24. Pulianmackal, L. T. et al. Multiple ParA/MinD ATPases coordinate the positioning of disparate cargos in a bacterial cell. Nat Commun 14, 3255 (2023).

25. Lee, P. S., Lin, D. C.-H., Moriya, S. & Grossman, A. D. Effects of the Chromosome Partitioning Protein Spo0J (ParB) on oriC Positioning and Replication Initiation in Bacillus subtilis. Journal of Bacteriology 185, 1326–1337 (2003).

26. Wang, X., Montero Llopis, P. & Rudner, D. Z. Bacillus subtilis chromosome organization oscillates between two distinct paàerns. Proceedings of the NaSonal Academy of Sciences 111, 12877–12882 (2014).

27. de Boer, P. A., Crossley, R. E. & Rothfield, L. I. Isolation and properties of minB, a complex genetic locus involved in correct placement of the division site in Escherichia coli. J Bacteriol 170, 2106–2112 (1988).

28. de Boer, P. A., Crossley, R. E. & Rothfield, L. I. A division inhibitor and a topological specificity factor coded for by the minicell locus determine proper placement of the division septum in E. coli. Cell 56, 641–649 (1989).

29. Hale, C. A., Meinhardt, H. & de Boer, P. A. Dynamic localization cycle of the cell division regulator MinE in Escherichia coli. EMBO J 20, 1563–1572 (2001).

30. Corbin, B. D., Yu, X.-C. & Margolin, W. Exploring intracellular space: function of the Min system in round-shaped Escherichia coli. EMBO J 21, 1998–2008 (2002).

31. Huang, K. C. & Wingreen, N. S. Min-protein oscillations in round bacteria. Phys. Biol. 1, 229 (2004).

32. Zieske, K. & Schwille, P. Reconstitution of self-organizing protein gradients as spatial cues in cell-free systems. Elife 3, e03949 (2014).

33. Ramirez-Arcos, S. et al. Deletion of the cell-division inhibitor MinC results in lysis of Neisseria gonorrhoeae. Microbiology 147, 225–237 (2001).

34. Ramirez-Arcos, S., Szeto, J., Dillon, J.-A. R. & Margolin, W. Conservation of dynamic localization among MinD and MinE orthologues: oscillation of Neisseria gonorrhoeae proteins in Escherichia coli. Mol Microbiol 46, 493–504 (2002).

35. Shaner, N. C. et al. A bright monomeric green fluorescent protein derived from Branchiostoma lanceolatum. Nat Methods 10, 407–409 (2013).

36. Gueiros-Filho, F. J. & Losick, R. A widely conserved bacterial cell division protein that promotes assembly of the tubulin-like protein FtsZ. Genes Dev 16, 2544–2556 (2002).

37. Jones, R. A., Yee, W. X., Mader, K., Tang, C. M. & Cehovin, A. Markerless gene editing in Neisseria gonorrhoeae. Microbiology 168, 001201 (2022).

38. Hsu, Y.-P. et al. Full color paleàe of fluorescent d-amino acids for in situ labeling of bacterial cell walls. Chem Sci 8, 6313–6321 (2017).

39. Stringer, C., Wang, T., Michaelos, M. & Pachitariu, M. Cellpose: a generalist algorithm for cellular segmentation. Nat Methods 18, 100–106 (2021).

40. Gadella, T. W. J. et al. mScarlet3: a brilliant and fast-maturing red fluorescent protein. Nat Methods 20, 541–545 (2023).

41. Pinho, M. G., Kjos, M. & Veening, J.-W. How to get (a)round: mechanisms controlling growth and division of coccoid bacteria. Nat Rev Microbiol 11, 601–614 (2013).

42. Veyrier, F. J., et al. Common Cell Shape Evolution of Two Nasopharyngeal Pathogens. PLOS GeneScs 11, e1005338 (2015).

43. Dalia, A. B. & Weiser, J. N. Minimization of bacterial size allows for complement evasion and is overcome by the agglutinating effect of antibody. Cell Host Microbe 10, 486–496 (2011).

44. Frirdich, E. et al. The Campylobacter jejuni helical to coccoid transition involves changes to peptidoglycan and the ability to elicit an immune response. Mol Microbiol 112, 280–301 (2019).

45. Dillard, J. P. Genetic Manipulation of Neisseria gonorrhoeae. Current Protocols in Microbiology 23, 4A.2.1-4A.2.24 (2011).

46. Grad, Y. H. et al. Genomic Epidemiology of Gonococcal Resistance to Extended-Spectrum Cephalosporins, Macrolides, and Fluoroquinolones in the United States, 2000-2013. J Infect Dis 214, 1579–1587 (2016).

47. Kohler, P. L., Hamilton, H. L., Cloud-Hansen, K. & Dillard, J. P. AtlA functions as a peptidoglycan lytic transglycosylase in the Neisseria gonorrhoeae type IV secretion system. J Bacteriol 189, 5421–5428 (2007).

48. Gunn, J. S. & Stein, D. C. Use of a non-selective transformation technique to construct a multiply restriction/modification-deficient mutant of Neisseria gonorrhoeae. Mol Gen Genet 251, 509–517 (1996).

49. Schindelin, J., et al. Fiji: an open-source plaäorm for biological-image analysis. Nat Methods 9, 676–682 (2012).

50. Ducret, A., Quardokus, E. M. & Brun, Y. V. MicrobeJ, a tool for high throughput bacterial cell detection and quantitative analysis. Nat Microbiol 1, 1–7 (2016).

51. Ershov, D. et al. TrackMate 7: integrating state-of-the-art segmentation algorithms into tracking pipelines. Nat Methods 19, 829–832 (2022).

