## Supplementary Figure 1 for "Axial asymmetry organizes division plane orthogonality in *Neisseria gonorrhoeae*"

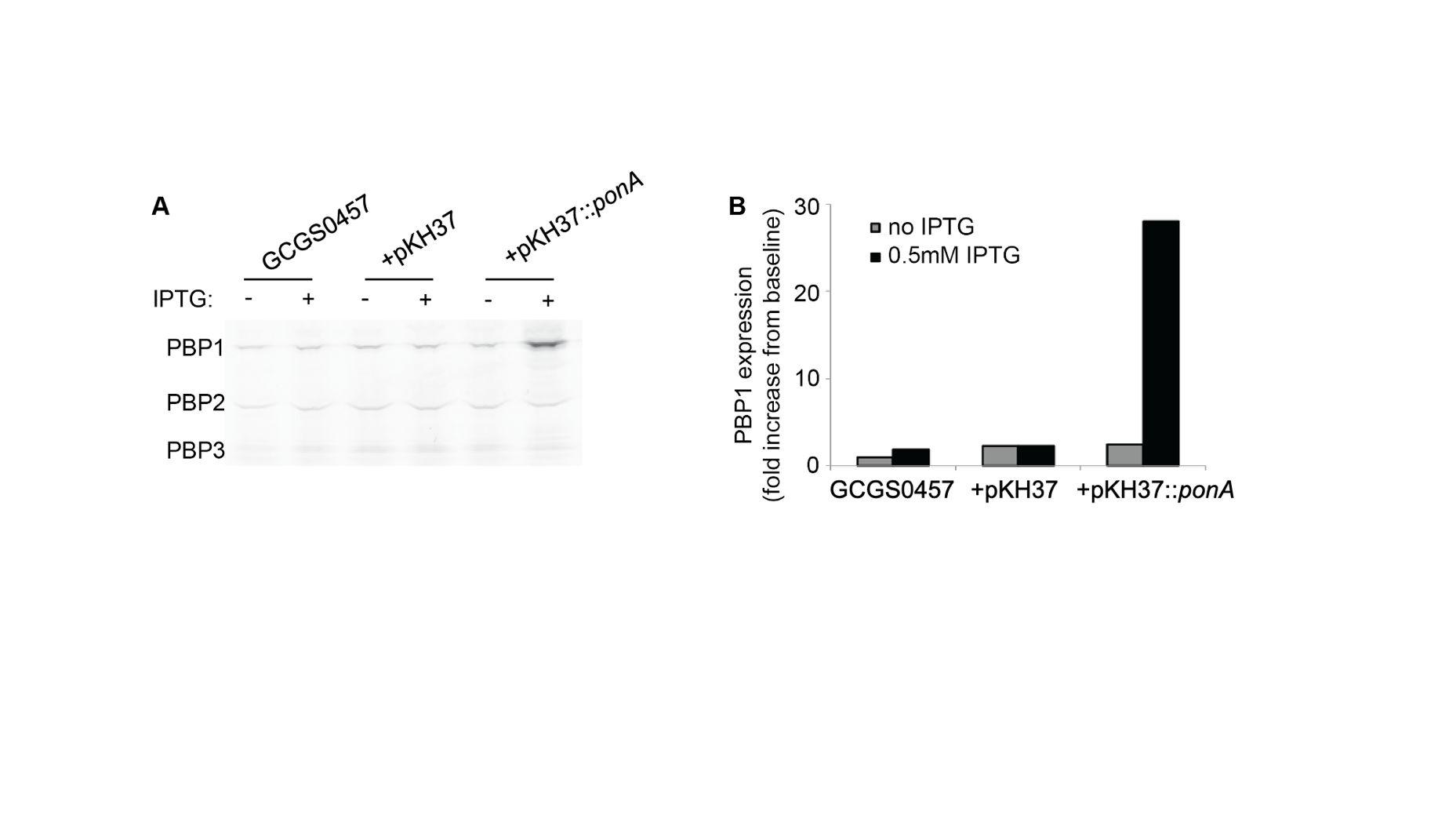

**Supplementary Figure 1. Overexpression of PBP1.** The parental strain GCGS0457 was transformed with the empty vector pKH37 or the PBP1 overexpression vector pKH37::*ponA*^L421P^. Each strain was grown in the presence or absence of 0.5mM IPTG to induce the lac promoter on pKH37 and stained with bocillin-FL to fluorescently label PBPs. Proteins were separated by SDS-PAGE. **(A)** Image of bocillin-FL fluorescence in the SDS-PAGE gel, visualized with the Typhoon FLA-9500 system. **(B)** PBP1 expression was quantified by densitometry and normalized to PBP1 expression in GCGS0457 without IPTG. When induced with 0.5mM IPTG,
the GCGS0457(pKH37::ponAL421P) strain produces approximately 30-fold more PBP1 than the
parental GCGS0457 strain

**Supplementary Video Legends**

**Supplementary Video 1, Supplementary Video 2, Supplementary Video 3.** All scale bars are 1 μm. Live cell imaging of nAB019. mNG-ZapA was imaged every 10 minutes.

**Supplementary Video 4, Supplementary Video 5.** All scale bars are 1 μm. Live cell imaging of nAB055. mScarlet3-ParB was imaged every 2 minutes, mNG-ZapA was imaged every 10 minutes. Red dots are ParB foci.
